# Elixirs of Life, threats to Human and Environmental Well-being: Assessment of Antibiotic Residues in Raw Meat Sold Within Central Market Kaduna Metropolis, Kaduna State, Nigeria

**DOI:** 10.1101/2022.01.04.474997

**Authors:** Habeeb Modupe Lateefat, Opasola Afolabi Olaniyi, Adiama Babatunde Yusuf, Ibrahim Azaman, Morufu Olalekan Raimi

## Abstract

Antibiotics, which are commonly used to treat human illnesses, are also used in animals for therapy, prophylaxis, and growth promotion. Sub-therapeutic antibiotic doses have typically been utilized for the last-mentioned purpose, which has contributed to resistance development. According to scientific data, certain antibiotic applications in food-producing animals can result in antibiotic resistance in intestinal bacteria, which can then be passed to the general population, causing treatment-resistant sickness. These antibiotic applications can also result in antibiotic resistance in non-pathogenic bacteria, whose resistance genes can be passed to disease-causing bacteria, resulting in antibiotic-resistant illnesses in people. Thus, this study assessed the antibiotics residues in raw meat sold in 6 slaughter houses in Kaduna State. The study is a descriptive cross-sectional study involving 6 slaughter houses in Central market Kaduna. Muscle, Kidney and liver samples were collected from each slaughterhouse. The antibiotic residues in the meat samples were analyzed using high performance liquid chromatography (HPLC) for tetracycline, ciprofloxacin and oxytetracycline residue results were presented in charts and tables. 18 different samples of beef (6 Muscles, 6 Liver and 6 Kidney) collected from abattoirs and meat vendors, the results shown that all beefs use three or more antimicrobial drugs. This research result revealed that 4(67%) tetracycline (oxytetracycline)were detected in meat samples at higher concentration), Oxytetraxycline (352.88 ± 221.58) of muscles is higher than (332.2± 217.05 of Liver and (263.33 ± 153.98) of Kidney is lower to muscles and liver. The Concentration of oxytetracycline were highest in muscles in samples 2. 3 and 6 which is above the WHO maximum residual limit. The concentration of streptomycin in the muscle, liver and kidney were detected (182.78 ± 56.23), (169.2 ± 58.39), (155.1 ± 50.20) but were within WHO Maximum residual limit. These high level of oxytetracycline residues in greater proportion of muscle samples destined for human consumption beyond MRLs could be as a result of the abuse of veterinary drugs as commonly practiced among livestock producers and vendors without observing withdrawal period prior to slaughter. The high-contamination rate of beef meat in the study areas is likely that consumers experience a high risk of exposure to drug residues.

## 1. Introduction

Chemical pollution was recognized as one of those limits whose continued impacts could degrade ecological and human resilience (Odipe *et al*., 2018; Suleiman *et al*., 2019; Omidiji and Raimi, 2019; Raimi *et al*., 2019; Raimi, 2019; Olalekan *et al*., 2020b; Olalekan *et al*., 2020c; Olalekan *et al*., 2020d; Adedoyin *et al*., 2020; Raimi *et al*., 2020b; Olalekan et al., 2021). As a result of the worldwide character of chemical pollution, a global response of internationally coordinated control measures is required, in addition to several local, regional, and national initiatives covering various groupings of compounds that are disconnected in time and place. The Stockholm Convention on Persistent Organic Pollutants (POPs) is one example of a global governance instrument, as it targets, at best, elimination, or, more broadly, sound management, of a collection of chemicals agreed upon through international talks. Chemical product chains, which span the life cycle stages from resource extraction to product manufacturing, usage, and disposal, are becoming increasingly complicated, frequently spanning many continents and decades, and offer new challenges to pollution control. For example, chemical production today can result in future emissions, particularly for chemicals in goods with long lifetimes.

While, antibiotics remain drugs of natural or synthetic origin that have the capacity to kill or to inhibit the growth of micro-organisms. Antibiotics that are sufficiently non-toxic to the host are used as chemotherapeutic agents in the treatment of infectious diseases of humans, animals and plants. Numerous researchers have reported that such chemical agents have been present in the environment for a long time, and have played a role in the battle between man and microbes (Isah *et al*., 2020a; Isah *et al*., 2020b; Olalekan *et al*., 2020; Raimi *et al*., 2020; Hussain *et al*., 2021a; Hussain *et al*., 2021b; Morufu *et al*., 2021; Morufu, 2021; Olalekan *et al*., 2021). They are chemically synthesized compounds that may reduce the proliferation and survival of different microorganisms (Demain & Sanchez, 2009; Raimi *et al*., 2020; Hussain *et al*., 2021a; Hussain *et al*., 2021b; Morufu *et al*., 2021; Morufu, 2021). Recently, antibiotics have emerge as broadly used in animal farming to protect and treat animals and to enhance growth promotion (Gelband *et al*., 2015). Antibiotic residues in food products of cattle’s, particularly meats, may prompt a negative impact, such as advancement of multidrug-resistant microbial strains (Chang *et al*., 2015; Kjeldgaard *et al*., 2012), allergic reactions and anaphylactic responses (Baynes *et al*., 2016), and the disturbance of typical digestive flora (Cotter *et al*., 2012). Also, resistant bacteria genes got from animal microbiomes might be evenly moved to human microbiota and to human pathogenic microscopic organisms (Marshall and Toll, 2011). Thus, the current study investigates the level of antibiotics residues in beef meat from selected abattoirs and meat vendors within Kaduna metropolis through determining the causes of antibiotic residues in the meat samples and assessing the possible risk of these antibiotic residues to consumers.

## 2. Materials and Methods

### Research Design

The research design is a cross-sectional descriptive survey where data collection will be at a single point in time from selected abattoirs and meat vendors within Kaduna metropolis.

### Sampling Location

The study was carried out within Kaduna state metropolis which is located between latitude 10.31’35.08N and longitude 7.26’19.64E (see figure 1 below).

**Figure 1.**
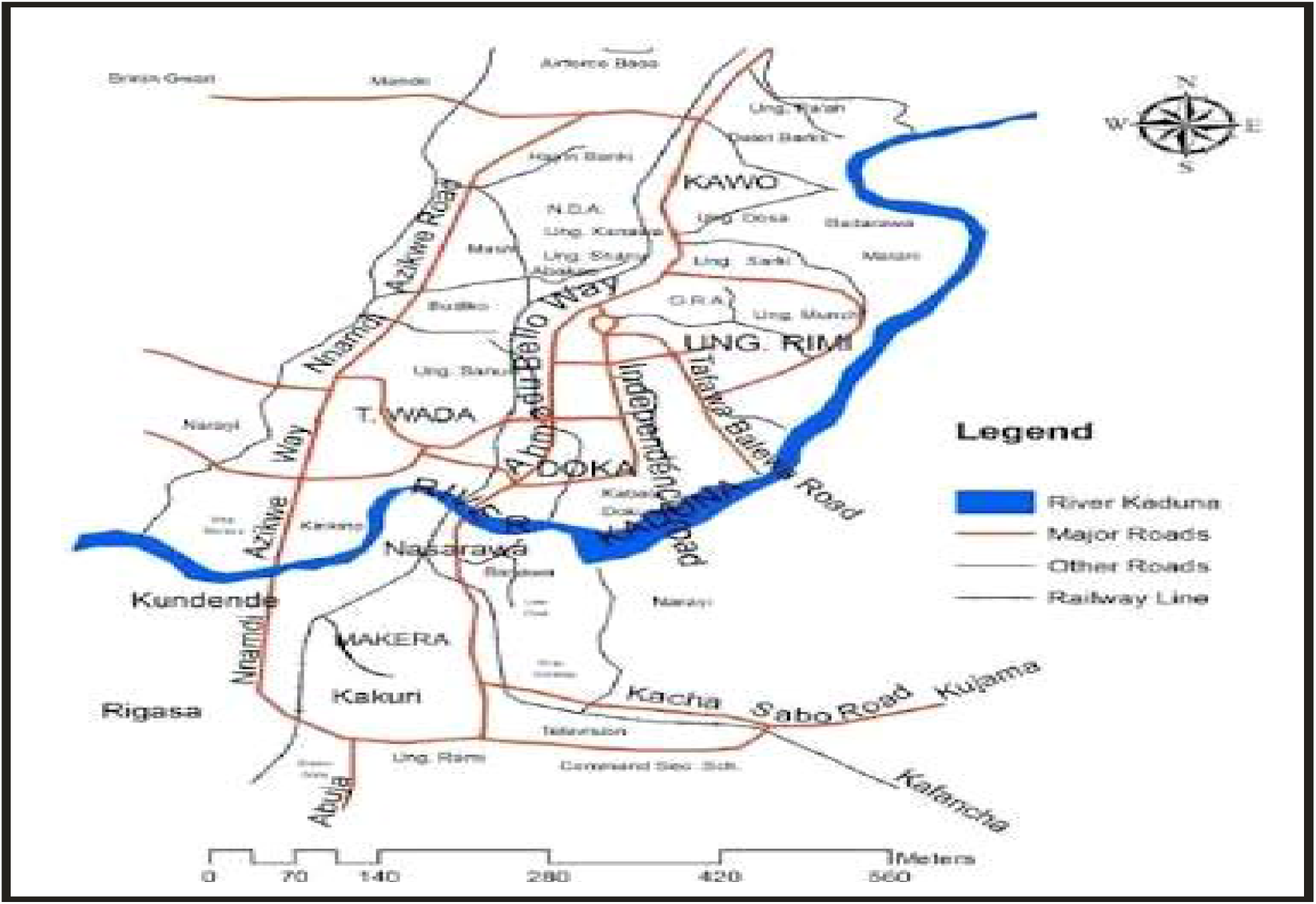
Map showing Kaduna metropolis.

### Sample Collection and Preparation

A total of 18 different samples (6 muscles 6 kidney and 6 liver) from beef meat was randomly collected from abattoirs and meat vendors within Kaduna metropolis. All samples were collected within recommended dates for consumption. After the collection, samples were packed in properly labeled sterile foil paper and transported to the laboratory for extraction.

### Apparatus

JASCO HPLC system with pump PU 580. A Rheodyric loop injector with injection volume of 20 U/L, JASO U.V defector 1575 operated at 565 am. JASO BORWIN software version 150 with Hercule 2000 chromatography interface for result integration and reordering, HiQSil, Ci8 L column (main column size: 4.6mm internal diameter x 75mm length and mobile phase made of 50m aqueous oxalic acid solution containing 13% (W/V) each of methanol and acetonitrile were used. Millipore all-glass filter unit comprising of glass funnel, holder, vacuum connector and receiving flask and vacuum flask with Barosiliate glass part in contact with liquid was used.

### Reagents

HPL water filtered through 0.2um with maximum impurities of 0.0003% and minimum transmission of 100% at 200 and 250nm produced from Qualigens fine chemicals, Glaxo smith wine pharmaceutical ltd was used. HPL grade acetonitrile and methanol purchased from Ranbaxy fine chemicals ltd, (SASNajur, india) as well as oxalic acid dehydrate (fluka, Acs, purity > 99.5%) and hydrochloric acid (fluka, HPCE grade filtered through a 0.22um membrane purchased from sigma-Aldrich cooperation) were used. Mobile phase was prepared and filtered through 0.22, Millipore Durapore solvent filters (disk 47mms, 9.6cm^3^ filtration area) under vacuum with Millipore all-glass filter unit degassed and delivered at a flow rate of 0.8 ml/min in isocratic mode.

### Data Analysis

The high-performance liquid chromatography (HPLC) relies on pumps to pass a pressurized liquid solvent containing the sample mixture through a column filled with a solid adsorbent material. Each component in the sample interacts slightly differently with the adsorbent material, causing different flow rates for the different components and leading to the separation of the components as they flow out the column. It has been applied for the detection of antimicrobials in meat, fish and internal organs (Kirbis *et al*., 2015).

### Statistical Analysis

The Statistical Analysis System (SAS software, 2010, Cary, NC, USA) (was used to analyzed the data obtained to present data in tables.

## 3. Results

The study in table 1 shows that (352.88 ± 221.58) of muscles is higher than (332.2 ± 217.05) of Liver and (263.33 ± 153.98) of Kidney is lower to muscles and liver.

**Table 1.**
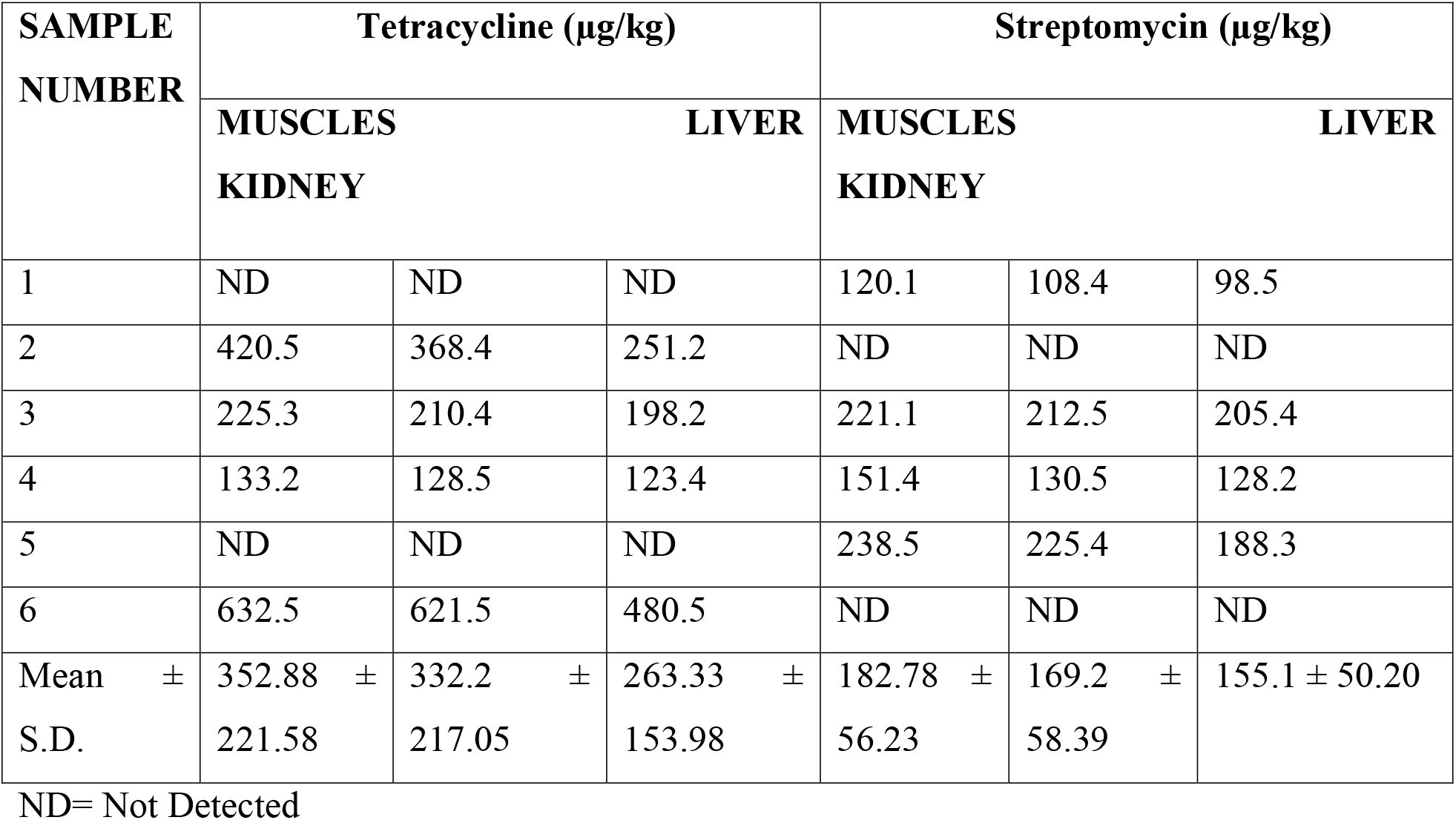
shows the residue levels of Tetracycline and Streptomycin in samples tissues.

### Antimicrobial drugs used in selected beef farms

A total of 18 different samples of beef (6 Muscles, 6 Liver and 6 Kidney) were collected from abattoirs and meat vendors the results shown that all beefs use three or more antimicrobial drugs. This research result revealed that almost all beef 12(201%) were using tetracycline (oxytetracycline). In addition to Streptomycin, 12(201%) and >MRLs 6(50%) (Table-2).

## 4. Discussion

From table 1 above, results revealed that Oxytetraxycline (352.88 ± 221.58) of muscles is higher than (332.2± 217.05 of Liver and (263.33 ± 153.98) of Kidney is lower to muscles and liver. The Concentration of oxytetracycline were highest in muscles in samples 2. 3 and 6 which above the WHO maximum residual limit (WHO,1997). The concentration of streptomycin in the muscle, liver and kidney were detected(182.78 ± 56.23), (169.2 ± 58.39), (155.1 ± 50.20) but were within WHO Maximum residual limit. The findings of this study was in line with the report of Elnasri and others (2018) who reported higher antibiotic residue in muscles (352.88 ± 221.58) than in the liver (332.2 ± 217.05), and kidney (263.33 ± 153.98). In contrast, Hala (2005) in Khartoum found an equal residue percentage of 24.6% in kidneys. This study was in contrast with the study that was conducted from October 2006 to May 2007 in Ethiopia to estimate the proportion of tetracycline levels in beef; the study focused on the Addis Ababa, Debre Zeit, and Nazareth slaughterhouses. Out of the total 384 samples analyzed for tetracycline residues, 71.3% had detectable oxytetracycline levels. Among the meat samples collected from the Addis Ababa, Debre Zeit, and Nazareth slaughterhouses, 93.8%, 37.5%, and 82.1% tested positive for oxytetracycline. The mean levels of oxytetracycline in muscle from the three slaughterhouses were as follows: Addis Ababa, 108.34 μg/kg; Nazareth, 64.85 μg/kg; and Debre Zeit, 15.916 μg/kg. Regarding kidney samples, oxytetracycline levels were found to be 99.02 μg/ kg in Addis Ababa, 109.35 μg/kg in Nazareth, and 112.53 μg/kg in Debre Zeit.

Form table 2 above 18 different samples of beef (6 Muscles, 6 Liver and 6 Kidney) collected from abattoirs and meat vendors the results shown that all beefs use three or more antimicrobial drugs. This research result revealed that 4(67%) tetracycline (oxytetracycline) were detected in meat samples at higher concentration above the WHO (1997) maximum residual limits and streptomycin were detectable in 4(67%) but are below the WHO (1997) MRLs. This study is comparable to a study in Tabriz, Iran, Abbasi *et al*., (2013) found 4(67%) of tetracycline residues in muscle samples, 4(67%) in liver samples, and 4(67%) in kidney samples in concentrations beyond the maximum residue limits (MRLs) of 6(50%). This study is also similar to a study carried out by Olufemi *et al*. 2009 in the Akure metropolitan abattoir were screened for oxytetracycline residues. Out of a total of 180 beef samples analyzed in this study, 98 (54.44%) had detectable levels of oxytetracycline of which 62 (34.44%) were beyond the MRLS by the WHO/FAO. In another report, analysis was carried out by Ibrahim *et al*. 2010 for 50 cattle slaughtered in the Sokoto metropolitan abattoir, Nigeria, in order to detect antibiotic residues in meat. A total of 44% of the slaughtered cattle tested positive. Penicillin was the drug with the highest rate of occurrence (14%) followed by tetracycline (8%) and Streptomycin. The high-contamination rate of beef meat in the study areas is likely that consumers experience a high risk of exposure to drug residues (Beyene *et al*., 2015).

**Table 2:**
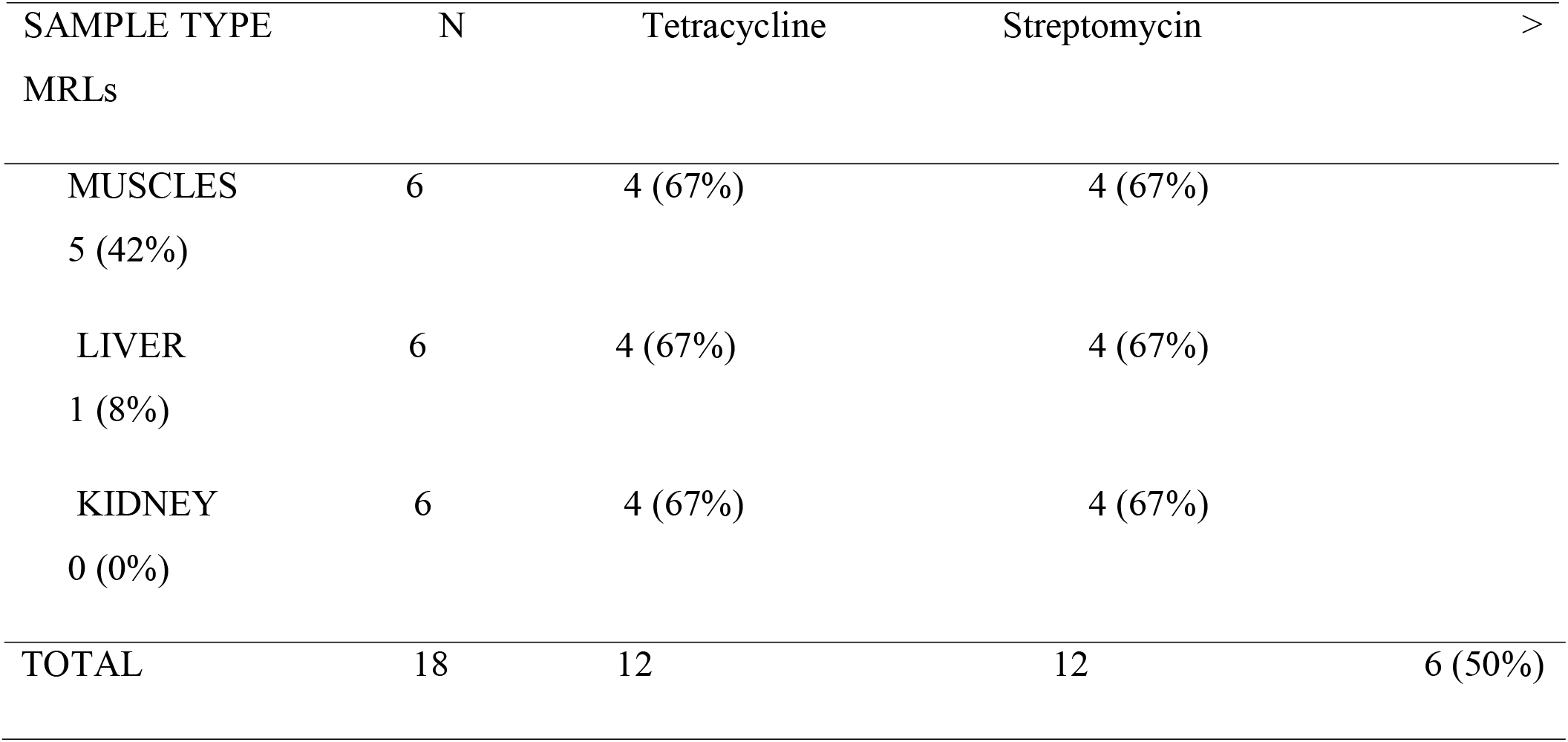
Percentage of positive samples with tetracycline and streptomycin.

## 5. Conclusion

As scientists, we must resist requesting greater scientific assurance before taking action, as this may postpone the implementation of control measures, which in this case will assist to halt widespread chemical pollution. Olalekan and others (Olalekan *et al*., 2020b; Olalekan *et al*., 2020d; Olalekan *et al*., 2020d; Raimi *et al*., 2020b; Morufu, 2021; Morufu *et al*., 2021f; Hussain *et al*., 21021a; Hussain *et al*., 2021b; Isah *et al*., 2020a; Isah *et al*., 2020b) have recorded cases where the call for further research to improve health risk assessments of chemicals resulted in action delays of up to several decades, despite early warnings of hazardous consequences. We must now go beyond Rachel Carson’s call to action on pesticides. With today’s phenomena of locally to globally distributed chemicals creating ill consequences, it is time to mobilize many people’s knowledge, capacity, and commitment to see Rachel Carson’s vision realized on a truly global scale. Thus, findings from this study have shown that the presence of veterinary drug residues in food products is not a health concern for Kaduna state alone but also, a global health concern as the consequence of using antimicrobial drugs to treat and prevent animal disease extend far beyond the farm. Therefore, the solution to antimicrobial residues will require a coordinated regulatory body to monitor the use of antimicrobial drugs to control diseases and also enforce punishment on indiscriminate usage. More so a sensitive, selective and reliable analytical method to easily detect and monitor antimicrobial residues in meat products should be encouraged.

## 6. Recommendations

Other measures that could be adopted to reduce antimicrobial residues in meat and meat products include:

1. Reduction of antimicrobial usage in livestock production (as many developed countries have banned its usage as growth promoters),
2. Enforcement of appropriate withdrawal periods of antimicrobial drugs application by government authorities or regulatory bodies before livestock slaughter.
3. Creation of awareness on implication of antimicrobial drugs residues in meat and meat products among consumers, and individual farmer,
4. Livestock producer should be educated on farm management, hygiene practices and antimicrobial usages in order to prevent occurrence of antimicrobial residues in meat production and lastly
5. Rapid screening methods should be developed for detecting and segregating samples containing above antimicrobial residues before the food products get to consumers. More so, establishment of framework to proper monitoring of drug usage and surveillance of antimicrobial resistance would be of great advantage.

## Notes

### Competing Interest Statement

The authors have declared no competing interest.

## References

Adedoyin OO, Olalekan RM, Olawale SH, et al (2020). A review of environmental, social and health impact assessment (Eshia) practice in Nigeria: a panacea for sustainable development and decision making. MOJ Public Health. 2020;9(3):81LJ87. DOI: 10.15406/mojph.2020.09.00328. https://medcraveonline.com/MOJPH/MOJPH-09-00328.pdf.

Baynes, RE., Dedonder, K., Kissell, L., Mzyk, D., Marmulak, T., Smith, G., Tell, L., Gehring, R., Davis, J., & Riviere, JE. (2016). Health concerns and management of select veterinary drug residues. Food and Chemical Toxicology, 88, 112–122. https://doi.org/10.1016/j.fct.2015.12.020.

Chang, Q., Wang, W., Regev-Yochay, G., Lipsitch, M., & Hanage, WP. (2015). Antibiotics in agriculture and the risk to human health: How worried should we be? Evolutionary Applications, 8(3), 240–247. https://doi.org/10.1111/eva.12185.

Demain, AL., & Sanchez, S. (2009). Microbial drug discovery: 80 years of progress. Journal of Antibiotics, 62(1), 5–16. https://doi.org/10.1038/ja.2008.16.

Gelband, H., Miller, PM., Pant, S., Gandra, S., Levinson, J., Barter, D., & Laxminarayan, R. (2015). The state of the world’s antibiotics 2015. Wound Healing Southern Africa, 8(2), 30–34. https://doi.org/10.10520/EJC180082

Hussain MI, Morufu OR, Henry OS (2021a). Probabilistic Assessment of Self-Reported Symptoms on Farmers Health: A Case Study in Kano State for Kura Local Government Area of Nigeria. Environmental Analysis & Ecology Studies 9(1). EAES. 000701. 2021. DOI: 10.31031/EAES.2021.09.000701. Pp. 975-985. ISSN: 2578-0336.

Hussain MI, Morufu OR, Henry OS (2021b) Patterns of Chemical Pesticide Use and Determinants of Self-Reported Symptoms on Farmers Health: A Case Study in Kano State for Kura Local Government Area of Nigeria. Research on World Agricultural Economy. Vol 2, No. 1. DOI: http://dx.doi.org/10.36956/rwae.v2i1.342. http://ojs.nassg.org/index.php/rwae/issue/view/31

Ibrahim, AI., Junaidu, AU and Garba, MK. (2010). Multiple antibiotic residues in meat from slaughtered cattle in Nigeria. Internet J. Vet. Med., 8: 1- DOI: 10.5580/1fcd.

Isah, HM., Sawyerr, HO., Raimi, MO., Bashir, BG., Haladu, S. & Odipe, OE. (2020a). Assessment of Commonly Used Pesticides and Frequency of Self-Reported Symptoms on Farmers Health in Kura, Kano State, Nigeria. Journal of Education and Learning Management (JELM), HolyKnight, vol. 1, 31–54. doi.org/10.46410/jelm.2020.1.1.05. https://holyknight.co.uk/journals/jelm-articles/.

Isah HM, Raimi MO, Sawyerr HO, Odipe OE, Bashir BG, Suleiman H (2020b) Qualitative Adverse Health Experience Associated with Pesticides Usage among Farmers from Kura, Kano State, Nigeria. Merit Research Journal of Medicine and Medical Sciences (ISSN: 2354-323X) Vol. 8(8) pp. 432–447, August, 2020. DOI: 10.5281/zenodo.4008682. https://meritresearchjournals.org/mms/content/2020/August/Isah%20et%20al.htm.

Kjeldgaard, J., Cohn, MT., Casey, PG., Hill, C., & Ingmer, H (2012). Residual antibiotics disrupt meat fermentation and increase risk of infection. MBio, 3(5), e00190.–e1112. https://doi.org/10.1128/mBio.00190-12.

Marshall, BM., & Levy, SB (2011). Food animals and antimicrobials: Impacts on human health. Clinical Microbiology Reviews, 24(4), 718 - 733. https://doi.org/10.1128/cmr.00002-11.

Morufu OR, Tonye VO, Ogah A, Henry AE, Abinotami WE (2021) Articulating the effect of Pesticides Use and Sustainable Development Goals (SDGs): The Science of Improving Lives through Decision Impacts. Research on World Agricultural Economy. Vol 2, No. 1. DOI: http://dx.doi.org/10.36956/rwae.v2i1.347. http://ojs.nassg.org/index.php/rwae/issue/view/31.

Morufu OR (2021). “Self-reported Symptoms on Farmers Health and Commonly Used Pesticides Related to Exposure in Kura, Kano State, Nigeria”. Annals of Community Medicine & Public Health. 1(1): 1002. http://www.remedypublications.com/open-access/self-reported-symptoms-on-farmers-health-and-commonly-used-pesticides-related-6595.pdf. http://www.remedypublications.com/annals-of-community-medicine-public-health-home.php.

Odipe OE, Raimi MO, Suleiman F (2018). Assessment of Heavy Metals in Effluent Water Discharges from Textile Industry and River Water at Close Proximity: A Comparison of Two Textile Industries from Funtua and Zaria, North Western Nigeria. Madridge Journal of Agriculture and Environmental Sciences. 2018; 1(1): 1–6. doi: 10.18689/mjaes-1000101. https://madridge.org/journal-of-agriculture-and-environmental-sciences/mjaes-1000101.php.

Olalekan MR, Abiola I, Ogah A, Dodeye EO (2021) Exploring How Human Activities Disturb the Balance of Biogeochemical Cycles: Evidence from the Carbon, Nitrogen and Hydrologic Cycles. Research on World Agricultural Economy. Volume 02, Issue 03. DOI: http://dx.doi.org/10.36956/rwae.v2i3.426. http://ojs.nassg.org/index.php/rwae.

Olalekan RM, Muhammad IH, Okoronkwo UL, Akopjubaro EH (2020). Assessment of safety practices and farmer’s behaviors adopted when handling pesticides in rural Kano state, Nigeria. Arts & Humanities Open Access Journal. 2020;4(5):191LJ201. DOI: 10.15406/ahoaj.2020.04.00170.

Olalekan RM, Oluwatoyin OA, Olawale SH, Emmanuel OO, Olalekan AZ (2020b) A Critical Review of Health Impact Assessment: Towards Strengthening the Knowledge of Decision Makers Understand Sustainable Development Goals in the Twenty-First Century: Necessity Today; Essentiality Tomorrow. Research and Advances: Environmental Sciences. 2020(1): 72–84. DOI: 10.33513/RAES/2001-13.https://ospopac.com/journal/environmental-sciences/early-online.

Olalekan RM, Dodeye EO, Efegbere HA, Odipe OE. Deinkuro NS, Babatunde A and Ochayi EO (2020c) Leaving No One Behind? Drinking-Water Challenge on the Rise in Niger Delta Region of Nigeria: A Review. Merit Research Journal of Environmental Science and Toxicology (ISSN: 2350-2266) Vol. 6(1): 031-049 DOI: 10.5281/zenodo.3779288

Olalekan RM, Oluwatoyin O and Olalekan A (2020d) Health Impact Assessment: A tool to Advance the Knowledge of Policy Makers Understand Sustainable Development Goals: A Review. ES Journal of Public Health; 1(1); 1002. https://escientificlibrary.com/public-health/in-press.php.

Olufemi, OI. and Ehinmowo AA (2009). Oxytetracycline Residues in Edible Tissues of Cattle Slaughtered in Akure, Nigeria. Internet J. Food Safety, 11: 62–66.

Omidiji AO and Raimi MO (2019) Practitioners Perspective of Environmental, Social and Health Impact Assessment (ESHIA) Practice in Nigeria: A Vital Instrument for Sustainable Development. Paper Presented at the Association for Environmental Impact Assessment of Nigeria (AEIAN) On Impact Assessment: A Tool for Achieving the Sustainable Development Goals in Nigeria, 7th and 8th November, 2019 In University of Port Harcourt.https://aeian.org/wp-content/uploads/2019/08/EIA-Presentations-Portharcourt.pdf.

Raimi MO, Sawyerr HO and Isah HM (2020) Health risk exposure to cypermethrin: A case study of kano state, Nigeria. Journal of Agriculture. 7th International Conference on Public Healthcare and Epidemiology. September 14-15, 2020 | Tokyo, Japan.

Raimi MO, Ihuoma BA, Esther OU, Abdulraheem AF, Opufou T, Deinkuro NS, Adebayo PA and Adeniji AO (2020b) “Health Impact Assessment: Expanding Public Policy Tools for Promoting Sustainable Development Goals (SDGs) in Nigeria”. EC Emergency Medicine and Critical Care 4.9 (2020).

Raimi MO, Omidiji AO, Adio ZO (2019) Health Impact Assessment: A Tool to Advance the Knowledge of Policy Makers Understand Sustainable Development Goals. Conference paper presented at the: Association for Environmental Impact Assessment of Nigeria (AEIAN) On Impact Assessment: A Tool for Achieving the Sustainable Development Goals in Nigeria, 7th and 8th November, 2019 in University of Port Harcourt. DOI: 10.13140/RG.2.2.35999.51366 https://www.researchgate.net/publication/337146101.

Raimi MO (2019) 21^st^ Century Emerging Issues in Pollution Control. 6^th^ Global Summit and Expo on Pollution Control May 06-07, 2019 Amsterdam, Netherlands.

Suleiman RM, Raimi MO and Sawyerr HO (2019) A Deep Dive into the Review of National Environmental Standards and Regulations Enforcement Agency (NESREA) Act. International Research Journal of Applied Sciences. pISSN: 2663-5577, eISSN: 2663- 5585. DOI No. Irjas.2019.123.123. www.scirange.com. https://scirange.com/abstract/irjas.2019.108.125.

